# RawBench: A Comprehensive Benchmarking Framework for Raw Nanopore Signal Analysis Techniques

**DOI:** 10.1101/2025.10.04.680405

**Authors:** Furkan Eris, Ulysse McConnell, Can Firtina, Onur Mutlu

## Abstract

Nanopore sequencing technologies continue to advance rapidly, offering critical benefits such as real-time analysis, the ability to sequence extremely long DNA fragments (up to millions of bases in a single read), and the option to selectively stop sequencing a molecule before completion. Traditionally, the raw electrical signals generated during sequencing are converted into DNA sequences through a process called *basecalling*, which typically relies on large neural network models. While accurate, these models are computationally intensive and often require high-end GPUs to process the vast volume of raw signal data. This presents a significant challenge for real-time processing, particularly on edge devices with limited computational resources, ultimately restricting the scalability and deployment of nanopore sequencing in resource-constrained settings. Raw signal analysis has emerged as a promising alternative to these resource-intensive approaches. While attempts have been made to benchmark conventional basecalling methods, existing evaluation frameworks 1) overlook raw signal analysis techniques, 2) lack the flexibility to accommodate new raw signal analysis tools easily, and 3) fail to include the latest improvements in nanopore datasets. Our goal is to provide an *extensible* benchmarking framework that enables designing and comparing new methods for raw signal analysis. To this end, we introduce RawBench, the *first* flexible framework for evaluating raw nanopore signal analysis techniques. RawBench provides modular evaluation of three core pipeline components: 1) reference genome encoding (using different pore models), 2) signal encoding (through various segmentation methods), and 3) representation matching (via different data structures). We *extensively* evaluate raw signal analysis techniques in terms of 1) quality and performance for read mapping, quality and performance for read classification, and 3) quality of raw signal analysis-assisted basecalling. Our evaluations show that raw signal analysis can achieve competitive quality while significantly reducing resource requirements, particularly in settings where real-time processing or edge deployment is necessary.

**CCS Concepts:** **Computing methodologies** → **Bioinformatics**; *Evaluation methodologies*; • **Applied computing** → **Computational genomics**.

**ACM Reference Format:** Furkan Eris, Ulysse McConnell, Can Firtina, and Onur Mutlu. 2025. RawBench: A Comprehensive Benchmarking Framework for Raw Nanopore Signal Analysis Techniques. In *Proceedings of the 16th ACM International Conference on Bioinformatics, Computational Biology, and Health Informatics (BCB ‘25), October 11–15, 2025, Philadelphia, PA, USA*. ACM, New York, NY, USA, 12 pages. https://doi.org/10.1145/3765612.3767302

## 1 Introduction

Nanopore sequencing [1–18] has revolutionized genomics by enabling the analysis of exceptionally long DNA molecules up to 4 million bases [19–22]. To sequence each base, nanopore devices measure ionic current changes as raw electrical signals while nucleic acids pass through nanoscale biological pores, called *nanopores* [4]. This approach offers two major benefits: 1) real-time sequencing decisions without having to fully sequence every read, a technique known as *adaptive sampling*, to reduce sequencing time and cost [23–30], and 2) natively providing richer information, such as epigenetic modifications (e.g., methylation patterns) [31–34].

There are two main approaches to analyzing nanopore sequencing data. The most widely adopted method is basecalling where raw electrical signals are translated into nucleotide sequences (i.e., A, C, G, T), called reads, using computationally intensive deep learning models [35–52]. This approach has the advantage of producing accurate reads that are compatible with existing bioinformatics pipelines. However, basecalling requires processing large segments of signal data [26], which increases latency and memory usage, and typically demands high-performance computing resources [21, 43, 53]. These requirements make real-time analysis challenging, i.e., processing and interpreting sequencing data as it is generated [26]. They also hinder portable sequencing, i.e., the use of field-deployable DNA/RNA sequencing devices such as Oxford Nanopore’s MinION, in resource-constrained environments [53]. While basecalling pipelines can be adapted for real-time use, doing BCB ‘25, October 11–15, 2025, Philadelphia, PA, USA so often requires reduction in quality and reliance on specialized or high-end hardware to meet latency constraints [54–61].

An emerging alternative that addresses these limitations is to analyze raw signals *directly* without basecalling [23–28, 53, 62–70]. By bypassing the translation step of electrical signals to text, direct analysis can significantly reduce computational overhead, lower memory usage, and enable faster decision-making. This is particularly advantageous in scenarios like adaptive sampling, where sequencing decisions must be made in real time [26].

Raw signal analysis offers unique opportunities to extract richer biological insights directly from raw data. Over the past years, a growing number of tools have been developed in this direction each advancing the state-of-the-art in terms of quality, speed, and resource usage [23–28, 53, 62–70]. To enable fair comparisons and better assessment of trade-offs, several benchmarking frameworks for raw nanopore signal analysis have been proposed [71, 72].s

However, existing benchmarking solutions lack three critical capabilities: (1) the ability to support both basecalled read and raw signal analysis (RSA), (2) the flexibility to incorporate newly developed methods targeting individual steps of raw signal analysis, and (3) access to standardized datasets that reflect the latest improvements in nanopore sequencing technology. These shortcomings hinder (1) *comprehensive* evaluation of raw signal analysis techniques, (2) high-resolution assessment that allows mixing and matching techniques from different tools, and (3) fair comparison of tools across different nanopore chemistries and raw signal behavior.

We identify three key challenges that underlie these shortcomings. First, prior benchmarking efforts [71, 72] adopt a task-focused approach, often focusing on a specific analysis goal, e.g., basecalling or modification detection, based on the capabilities of a small number of tools. This limits broader insight, as the chosen task may not reflect the generalizable strengths of different raw signal analysis techniques. Second, prior benchmarking solutions are designed as monolithic systems to ease implementation, and hence the modularity needed to flexibly incorporate and evaluate new methods. Third, there is limited availability of comprehensive and standardized raw signal datasets, due to both the default discarding of the raw nanopore signal data during basecalling (e.g., by ONT’s MinKNOW software) and the general lack of public sharing [73].

**Our goal** is to enable comprehensive and extensible benchmarking of raw nanopore signal analysis methods that support both basecalled and RSA approaches on large, representative datasets. To this end, we propose RawBench, the *first* modular raw nanopore signal analysis benchmarking framework designed to address key limitations of existing benchmarking frameworks.

We demonstrate the capabilities of RawBench through an extensive benchmarking study, evaluating two commonly used tasks, i.e., read mapping and read classification, that can leverage raw signal analysis. Our study systematically evaluates thirty diverse algorithmic strategies across a wide range of four datasets and two experimental conditions. Our key contributions include:

- We benchmark 30 unique method combinations for raw signal analysis, exploring key design trade-offs and scaling characteristics across tasks and sequencing conditions. Our comprehensive structure provides insights critical for realtime use and resource-constrained environments.
- We enable flexible integration of new raw signal analysis methods, facilitating fine-grained comparisons and highresolution method development, addressing the inflexibility of prior benchmarking tools and establishing the infrastructure necessary for well-documented and open progress in raw signal analysis.
- We incorporate curated, up-to-date raw nanopore datasets spanning multiple species and sequencing chemistries, providing a robust foundation for fair and reproducible evaluations.
- We fully open source RawBench’s codebase, datasets, and benchmark results to foster transparency and accelerate future research in raw nanopore signal analysis. 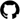

## 2 Background, Related Work, Our Goal

Nanopore sequencing uniquely enables *adaptive sampling*, a technique that decides in real time whether to continue or stop reading a DNA molecule based on its initial signal reading. This lets the system focus on molecules that are likely to be important (such as genes linked to a disease) while skipping irrelevant ones. Unlike traditional enrichment methods that require special lab preparation, adaptive sampling performs this selection during sequencing, saving time and cost [29, 74]. When combined with the portability of nanopore sequencers, adaptive sampling enables field applications such as rapid pathogen surveillance [75, 76]. However, real-time decision-making aspect of adaptive sampling imposes stringent latency constraints, as decisions must occur fast enough to fully utilize pore throughput [26, 64], demanding lightweight methods on resource-limited edge devices [53].

Basecalling is the dominant approach to analyzing nanopore signals, where deep neural networks (e.g., convolutional / recurrent architectures) translate raw ionic current signals into basecalled nucleotide sequences (i.e., A, C, G, T) [35–52]. State-of-the-art basecallers like Dorado [40] achieve high quality but rely on complex, resource-intensive architectures that create bottlenecks for downstream genomic analyses. This overhead arises from processing long signal traces, often millions of data points per read, using neural networks with substantial redundancy, as evidenced by a recent work showing that up to 85% of model weights in such models can be pruned without significant quality loss [43]. Consequently, basecalling bottlenecks real-time portable sequencing [25, 53, 59], motivating efforts to accelerate performance via hardware and algorithmic optimizations [77–79].

To accelerate basecalling, several works explore FPGAs [43, 54– 56, 80, 81] and PIM-based solutions [57–60], leveraging parallelism and eliminating data movement overhead to reduce latency and energy consumption. GenPIP [60] concurrently executes basecalling and read mapping, the process of aligning sequenced reads to a reference genome, inside memory to minimize data movement and redundant computation. CiMBA [59] advances the field with a compact compute-in-memory accelerator and analog-aware networks for on-device basecalling. While effective, these solutions require specialized hardware, limiting their adaptability to changes in the sequencing technology and deployment scenario.

A promising alternative to basecalling is RSA, which operates directly on electrical signals without translating them into reads [26, 64, 65]. These techniques typically involve three distinct stages. (1) In the *reference encoding* stage, a genome sequence is converted into expected corresponding electrical signals using a pore model that predicts current levels for each DNA k-mer, i.e., short nucleotide sequences of length *k*. (2) In the *signal segmentation* stage, the continuous nanopore current is divided into discrete segments corresponding to individual k-mers. This segmentation step explicitly extracts meaningful segments from noisy raw signal data using statistical or learning methods, unlike basecalling where segmentation is performed implicitly in neural networks. (3) In the *representation matching* stage, the segmented signals are compared against the encoded reference using different matching algorithms to find similarities despite the noise in the signal data. More details on methods addressing each of these stages can be found in Section 3.

Beyond algorithmic developments for RSA, recent works demonstrate the benefits of real-time raw signal processing on specialized hardware. SquiggleFilter [25] uses an ASIC to filter irrelevant reads before basecalling, eliminating unnecessary and costly basecalling and enabling fast pathogen detection. HARU [53] introduces an FPGA-based accelerator for real-time adaptive sampling in resource-constrained environments. MARS [61] adopts a storage-centric Processing-In-Memory (PIM) approach, i.e., where computation is performed directly inside or near memory, to accelerate raw signal genome analysis, combining filtering, quantization, and in-storage execution to achieve up to 28× speedup and 180× energy savings over software baselines. Together, these efforts underscore the growing interest in hardware-software co-design for raw signal analysis and unlock capabilities for the diverse and dynamic landscape of nanopore sequencing applications.

As RSA grows, particularly in latency- and energy-sensitive settings, there is an increasing need for fair, systematic evaluation of emerging tools and techniques. Benchmarking becomes critical not only to measure performance but also to understand the trade-offs between different design choices. However, existing benchmarking efforts have limitations. While NanoBaseLib [71] and basecalling benchmarks [72] provide valuable foundations, they have a fundamental limitation: they completely disregard RSA tools. This gap is compounded by other issues, including a lack of biological justification for design choices (e.g., training data generation), limited adaptability to new chemistries, and benchmarking of tools as a whole that obscures component-level contributions.

Beyond the software benchmarks for genomic analysis, several benchmarking frameworks have emerged to evaluate the performance of different genomic analysis methods on hardware platforms. Genomics-GPU [82] offers GPU-accelerated workloads for genome comparison, matching, and clustering. GenomicsBench [83] covers data-parallel kernels for short- and long-read workflows on CPU/GPU. While these frameworks contribute to systematic hardware-software co-design for genomics, they do not focus on the unique challenges of RSA.

**Our goal** is to introduce an extensible benchmarking framework designed to systematically evaluate and compare text-based and RSA methods. RawBench enables: 1) a more inclusive setting towards RSA; 2) modular assessment of different reference encoding, signal encoding, and representation matching techniques; and 3) compatibility with diverse sequencing chemistries and organisms. By enabling fair, component-level evaluation, RawBench helps and accelerates the development of biologically informed and computationally efficient raw nanopore signal analysis methods.

## 3 RawBench Framework

RawBench is a benchmarking framework for raw signal analysis (RSA) to evaluate RSA methods across nanopore chemistries, datasets, and computational settings, based on ground truth generated using expensive basecalled analysis. The framework is structured into three RSA stages as shown in Figure 1: ➊ encoding the reference genome into expected signal patterns, ➋ segmenting raw signals into a comparable encoded representation, and ➌ matching these two encoded representations for tasks such as read mapping or classification.

**Figure 1.**
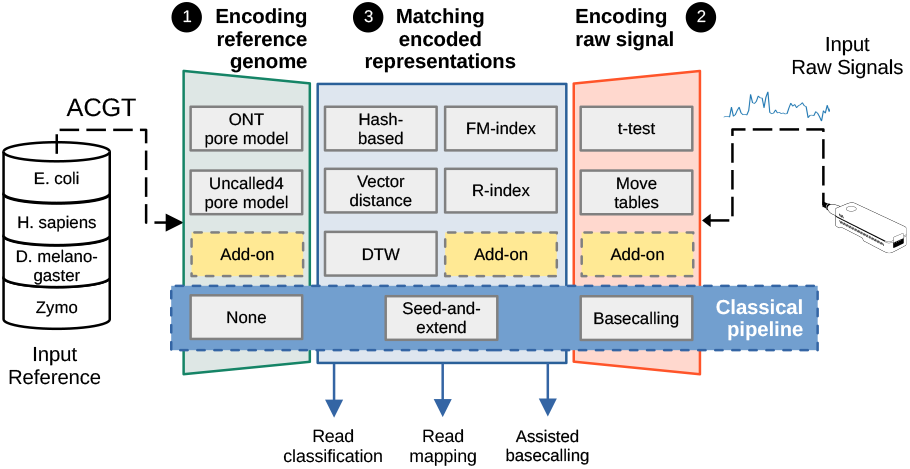
Overview of RawBench.

The framework’s modular design enables as inputs (i) reference genomes, (ii) nanopore raw signals from multiple chemistries and organisms, and (iii) any RSA method that targets one or more of the three RawBench stages. To support comprehensive evaluation, RawBench includes datasets spanning bacteria, eukaryotes, and metagenomes from different nanopore chemistries, while also allowing the community to easily integrate their own datasets or downstream tasks. Beyond quality, RawBench also reports runtime and memory, providing practical insights for deploying RSA methods in real-world sequencing workflows.

### 3.1 Encoding the Reference Genome

To enable the alignment of raw signals to their genomic origins directly in raw signal space, the reference genome must first be translated into a sequence of expected electrical signal patterns. This is typically done using k-mer models, which map k-mers to their characteristic electrical signal distributions. However, these models are tightly coupled to specific nanopore chemistries and flow cell versions, limiting generalizability. Tools like Uncalled4 [84] and Poregen [85] address this problem by learning k-mer models *de novo* which enables faster adaptation to new chemistries or settings.

Despite their central role in RSA, the effectiveness of different k-mer models remains poorly understood with limited systematic benchmarking across chemistries. Factors critical for field deployment decisions such as accuracy, memory, and runtime can vary unpredictably depending on the pore model, underscoring the need for robust evaluation.

RawBench contributes by providing a unified framework to systematically benchmark k-mer models across multiple nanopore chemistries. It enables fair comparison of both official ONT and open-source learned models, measuring not only the downstream task quality but also memory and runtime trade-offs. In doing so, RawBench enables accurate evaluation of how reference encoding choices impact RSA quality and performance, guiding informed deployment decisions in field settings.

### 3.2 Segmenting the Raw Signals

To enable accurate alignment between raw signals and the reference genome, raw signals must be transformed into latent representations that meaningfully correspond to nucleotide sequences. This transformation step is central to downstream tasks such as read mapping and classification, which involves determining the origin or type of a DNA/RNA read, such as its species, gene, or functional category. Two dominant strategies exist for this transformation.

First, RSA requires explicit segmentation of raw signals into discrete segments corresponding to k-mers. Segmentation methods include t-test changepoint detection [86] and resquiggling [68, 87], though these often lack awareness of sequencing context or pore-specific characteristics, limiting robustness across nanopore chemistries. A context-aware alternative is Campolina [88], a convolutional neural network (CNN) trained to predict segmentation points from raw signals. Unlike statistical methods such as t-test, which operate only on local fluctuations within a signal chunk, Campolina leverages broader sequencing and chemistry priors to improve robustness at the cost of requiring extensive pretraining.

The second approach, basecalling, employs neural networks with CRF [89] and CTC [90] decoders to implicitly convert signals to nucleotides. Basecallers process overlapping signal chunks that are concatenated post-processing, benefitting from a relatively long context compared to RSA approaches. Move tables, an intermediate output extracted from basecallers, can provide coarse segmentation points for RSA [85]. Only Bonito [91] and Dorado [40] basecallers support the latest R10.4.1 chemistry, but rely on proprietary training data, limiting adaptability and reproducibility.

RawBench enables systematic evaluation of segmentation approaches by providing a wide range of benchmarks across chemistries and sequencing contexts, allowing fair comparison of statistical and learning-based methods. By exposing the contrasting characteristics of different segmentation approaches, RawBench helps identify opportunities for developing future segmentation methods, e.g., those that combine the efficiency of statistical techniques with the robustness of learning-based models.

### 3.3 Matching Encoded Representations

To complete the mapping process from raw signals to reference genomes, the encoded representations of both signals and reference must be matched in a way that preserves biological accuracy while maintaining computational efficiency. Due to the high dimensionality and variability of raw signal data, designing scalable and accurate matching algorithms remains a significant challenge.

Matching approaches generally fall into two main categories based on their computational demands and degrees of sensitivity: (1) heuristic methods that prioritize speed and scalability, often using hashing or indexing to perform approximate matching; and (2) precise alignment methods that emphasize quality, typically leveraging distance metrics or dynamic programming. Each approach offers different trade-offs in terms of speed, memory, and downstream utility, especially when handling complex genomes and high-throughput raw signal streams. RawBench enables side-by-side evaluation of these techniques under thirty realistic workloads.

The first category includes hash-based and probabilistic methods. RawHash [64] and RawHash2 [65] use quantization to map similar raw signal segments to shared hash buckets, enabling fast approximate matching for mapping against large genomes. UNCALLED [27] employs a probabilistic FM-index to estimate the likelihood of signal-to-k-mer matches. Sigmap [26] embeds both signal and reference into a shared high-dimensional space, but suffers from computational overhead and the curse of dimensionality [92]. Compressed indexing strategies like the R-index [93, 94] offer an alternative, supporting lightweight exact matching over repetitive regions. Tools like Sigmoni [66] use this structure to compute pseudo-matching lengths (PMLs), where longer PMLs indicate stronger matches between compressed signal and reference segments.

The second category focuses on fine-grained alignment methods, most notably Dynamic Time Warping (DTW) [95, 96], which has been employed in earlier works to achieve high-quality alignments between raw signals and reference sequences [25, 53, 68]. While DTW offers precise signal-to-reference alignment, its computational cost is prohibitive with larger genomes, especially in real-time scenarios. To mitigate this, hybrid approaches such as RawAlign [63] combine fast seeding techniques [64, 65] with selective DTW refinement, achieving a practical trade-off between quality and performance.

RawBench evaluates both types of strategies across varying genome complexities and computational demands. This extensive framework encourages rapid development and comprehensive evaluation of raw signal-to-reference matching algorithms.

### 3.4 Assisting Basecalled Analysis

To improve efficiency in conventional basecalling pipelines, it is often desirable to avoid unnecessary basecalling of reads that are unlikely to contribute to downstream analyses. The goal of pre-basecalling raw signal filtering, i.e., the process of inferring raw signals that are likely to be important from signal characteristics, is to identify such reads directly from raw signals and forward only promising reads to the basecaller.

By default, RSA tools process a prefix of raw signals and stop once a mapping is found to meet the constraints of real-time analysis, limiting the amount of surrounding signal available for highly precise downstream analyses. Pre-basecalling raw signal filtering transforms RSA into a computationally efficient preprocessing step. By filtering out relatively insignificant raw signals before full basecalling, it is possible to reduce the load on basecallers, conserve computational resources, and improve overall throughput. Filtering is performed independently of the basecaller, making it compatible with existing pipelines without altering downstream aspects.

Tools like TargetCall [35] showcase the benefits of pre-basecalling filtering by performing an initial lightweight raw signal analysis, allowing for more accurate and resource-intensive basecallers to focus on important reads. Within RawBench, such pre-filtering approaches can be evaluated systematically across different datasets and signal complexities, enabling assessment of trade-offs between computational savings and downstream analysis quality. This makes RawBench a powerful tool to guide the design and deployment of future pre-basecalling strategies in practical sequencing workflows.

## 4 RawBench Datasets

To evaluate the robustness across varying genomic complexities, RawBench includes datasets from *E. coli, D. melanogaster, H. sapiens*, and a *Zymo* metagenomic dataset as shown in Table 1. Dataset names in Table 1 are provided as hypertexts referring to the source and preprocessing scripts are provided in.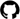

**Table 1:** Summary of RawBench datasets.

| Dataset | Genome Size (Mbp) | Number of Reads | Number of Bases (Mbp) | Depth of Coverage |
| --- | --- | --- | --- | --- |
| E. coli | 4.6 | 180,000 | 1,131 | 225× |
| D. melanogaster | 143.7 | 231,278 | 1,730 | 12.04× |
| H. sapiens | 3,200 | 16,000 | 329 | 0.11× |
| Zymo mock | 65.4 | 10,000 | 134 | 2.05× |

We use nanopore sequencing data generated using the old R9.4.1 and the latest R10.4.1 chemistry. All datasets provide raw signals (i.e., FAST5 or POD5) and basecalled reads (i.e., FASTQ). We note that alternative formats like SLOW5 improve storage efficiency and read performance [97]. When possible, we prefer 400 bps (i.e., bases per second passing through the nanopore) mode over the deprecated 260 bps for better quality and higher sequencing yield [40].

Each dataset provides sufficient coverage for downstream tasks such as single nucleotide polymorphism (SNP) and structural variant (SV) calling [98–101]. Summary statistics are shown in Table 1.

RawBench spans simple to complex genomes for evaluating scalability of different RSA techniques. *E. coli* represents a compact bacterial genome for rapid testing. *D. melanogaster* adds moderate complexity with repetitive and heterochromatic regions. *H. sapiens* provides a highly complex genome rich in SVs, repeats (>45%), and allelic diversity, posing the most demanding read mapping challenge. The *Zymo* mock community introduces a metagenomic case for mixed-species classification. Datasets originating from reference sequences with contrasting characteristics ensures that benchmarked RSA techniques remain robust across diverse raw nanopore sequencing data analyses.

## 5 Evaluation

### 5.1 Evaluation Methodology

We implement RawBench as a modular Nextflow [102] framework with C++ components, enabling plug-and-play RSA stages from Section 3. In particular, stage-level techniques (e.g., t-test–based segmentation and hash-based matching) are provided as standalone C++ modules and they can be mixed and matched. For completeness, we also provide benchmarking scripts that invoke prebuilt binaries of established RSA tools; while these wrappers facilitate fair, outof-the-box comparisons, they naturally limit full combinatorial exploration compared to our C++ modules.

#### Methods

We evaluate quality, performance and coverage of different RSA techniques. To assess read mapping and classification quality, we create thirty different RSA pipeline combinations out of the RawBench component pool. Outputs are compared against ground truth generated by the Dorado [40] super-accurate (SUP) basecaller followed by the minimap2 [103] read mapper. Metrics include true positives (TP, i.e., correctly mapped reads), false positives (FP, i.e., incorrectly mapped reads), false negatives (FN, i.e., reads that could not be mapped), not aligned (NA, i.e., reads with no ground truth location), precision 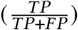, recall 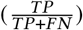, and F1 score 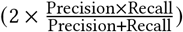. Results are computed using UNCALLED pafstats [27].

For performance, we report elapsed time (wall-clock time), CPU time, and peak memory usage. GPU acceleration is used only for Dorado, while all RSA components (pore models, segmentation, matching) run on CPU to ensure fair comparison and reflect typical deployment scenarios.

We benchmark end-to-end basecalled read and raw signal analysis while considering the impact of (1) pore models, (2) segmentation methods, (3) matching methods, and (4) RSA-assisted basecalling and 5) basecaller context length.

Unless otherwise stated, evaluations fix the pore model to Uncalled4 [84], segmentation to t-test [86], and matching to hash-based [104] (for read mapping and RSA-assisted basecalling) or R-index [93, 94] (for read classification).

#### Pore Models

First, we evaluate how well different *pore models* translate DNA into expected signal features. We isolate the effect of two pore models, ONT [40] and Uncalled4 [84], on downstream quality and performance, corresponding to stage ➊ (Fig. 1).

#### Segmentation Methods

Second, we assess three *segmentation* strategies based on their ability to partition raw signals into biologically meaningful events: 1) t-test [86], 2) move tables [40], and 3) neural network-based method Campolina [88], corresponding to stage ➋ (Fig. 1).

#### Matching Methods

Third, we compare five different *matching* algorithms to determine their effectiveness in signal-to-reference alignment: 1) hash-based [104], 2) FM-index [105], 3) vector distances [26], 4) R-index [93, 94], and 5) DTW [95, 96], corresponding to stage ➌ (Fig. 1).

#### Assisting Basecalled Analysis

Fourth, we evaluate RSA as a pre-filtering step [25, 35, 53] for basecalling by comparing depth of coverage with and without limiting Dorado’s context to successfully mapped reads from RSA.

Fifth, we evaluate the effect of limiting Dorado’s context to 1, 2, 5, or 10 chunks of 4000 signal points each, corresponding to approximately 1 second of data [70]. This reveals insights about the quality–performance trade-offs by adjusting context length.

#### Experimental Setup

We perform all experiments on a server equipped with an NVIDIA A6000 GPU [106] and an Intel(R) Xeon(R) Gold 6226R CPU [107] running at 2.90 GHz. We conduct each evaluation with 64 threads. We show the parameters and versions of the tools we evaluate in Supplementary Tables S7 and S8 respectively.

### 5.2 Quality Evaluation

Tables 2 and S1 show the quality of two pore models: ONT and Uncalled4 when applied across datasets and downstream tasks. We make two key observations.

**Table 2:** Read mapping quality using different pore models.

| Pore Model | F1 | Precision | Recall |
| --- | --- | --- | --- |
| *E. coli* |  |  |  |
| ONT | 0.79 | 0.88 | 0.71 |
| Uncalled4 | **0.83** | **0.91** | **0.77** |
| *D. melanogaster* |  |  |  |
| ONT | 0.66 | 0.93 | 0.51 |
| Uncalled4 | **0.72** | **0.94** | **0.59** |
| *H. sapiens* |  |  |  |
| ONT | 0.58 | 0.86 | 0.44 |
| Uncalled4 | **0.66** | **0.87** | **0.53** |

First, pore model choice has a noticeable impact on downstream analysis quality, with Uncalled4 outperforming ONT on read mapping tasks across all organisms (see Table 2). Uncalled4 achieves higher F1 scores (e.g., 0.83 for *E. coli*), driven largely by improvements in recall while maintaining high precision (≥ 0.87). In contrast, ONT pore models consistently show lower recall, particularly in larger genomes (0.44 for *H. sapiens*), limiting their ability to provide true alignments. These results concur with prior findings [84], which highlight limitations in ONT pore modeling for R10.4.1 chemistry, motivating more expressive reference-to-signal representations.

Second, the benefits of Uncalled4 appear to diminish in the R9.4.1 read classification task (see Table S1). Uncalled4 and ONT pore models provide identical quality. This observation is in line with previous work [27], which shows that ONT and Uncalled4 pore models for the old R9.4.1 chemistry exhibited almost identical characteristics. This suggests that while models like Uncalled4 can substantially improve mapping performance, they do not necessarily translate into comparable gains in higher-level tasks.

Together, these findings indicate that pore models remain an important determinant of read mapping quality. While advances such as Uncalled4 demonstrate clear improvements over ONT in read mapping, read classification results suggest that further research is needed to develop pore models that generalize their benefits across different downstream tasks and nanopore chemistries.

#### Segmentation Methods

Table 3 shows the quality of three segmentation approaches: t-test, move tables, and Campolina when applied to each task and dataset. We make two key observations. First, segmentation quality has a substantial impact on downstream task quality, with Campolina outperforming the other two approaches significantly on read mapping tasks, in particular for the larger *H. sapiens* genome (F1 = 0.79). t-test segmentation outperforms move tables across all organisms. t-test achieves high mapping F1 scores (up to 0.83 for *E. coli*) and excels in classification (F1 = 0.92 on *Zymo*), maintaining strong precision (*>*= 0.85) across all tasks. In contrast, move tables yields significantly lower quality, likely due to its coarse, stride-based segmentation that introduces noise and fails to capture many signal transitions.

**Table 3:** Read mapping quality using different segmentation methods.

| Segmentation Method | F1 | Precision | Recall |
| --- | --- | --- | --- |
| *E. coli* |  |  |  |
| t-test | 0.83 | 0.91 | 0.77 |
| Move tables | 0.07 | 0.07 | 0.06 |
| Campolina | **0.89** | **0.94** | **0.85** |
| *D. melanogaster* |  |  |  |
| t-test | **0.72** | 0.94 | **0.59** |
| Move tables | 0.05 | 0.23 | 0.03 |
| Campolina | **0.72** | **0.95** | 0.57 |
| *H. sapiens* |  |  |  |
| t-test | 0.66 | 0.87 | 0.53 |
| Move tables | 0.01 | 0.11 | 0.01 |
| Campolina | **0.79** | **0.96** | **0.67** |

Second, genomic complexity negatively affects recall, even for high-performing methods like t-test and Campolina. While precision remains mostly stable, recall drops with larger genomes, reflecting the challenge of accurately segmenting signals from complex genomes. This observation also holds for the neural network-based approach Campolina, albeit with some improvements in recall for the *H. sapiens* data, pointing to unsolved underlying challenges in segmentation. This trend highlights a need for segmentation approaches that retain high precision while improving sensitivity to capture true segments in raw signals.

These findings show that segmentation remains a critical bottleneck: methods like t-test provide strong precision but falter on recall in complex genomes, pointing to the need for approaches that can maintain sensitivity without sacrificing precision.

#### Matching Methods

Tables 4 and 5 show the quality of five matching approaches: hash-based methods, FM-index, vector distances, R-index and Dynamic Time Warping (DTW). We note that the combination of DTW and t-test segmentation forms the basis of the f5c resquiggle method [68]. We make five key observations. First, hash-based matching provides consistently high quality across mapping tasks, achieving high F1 scores across organisms. This approach, used in tools like RawHash and RawHash2 [64, 65], excels through its ability to quickly identify approximate matches using locality-sensitive hashing, making it computationally efficient for large-scale analyses while maintaining high quality.

**Table 4:** Read mapping quality using different matching techniques.

| Matching Method | F1 | Precision | Recall |
| --- | --- | --- | --- |
| *E. coli* |  |  |  |
| Hash-based | 0.83 | 0.91 | 0.77 |
| FM-index | 0.23 | 0.13 | 0.80 |
| Vector distances | 0.83 | 0.84 | **0.82** |
| R-index | 0.67 | 0.79 | 0.58 |
| DTW | **0.86** | **0.99** | 0.75 |
| *D. melanogaster* |  |  |  |
| Hash-based | 0.72 | 0.94 | 0.59 |
| FM-index | 0.02 | 0.17 | 0.01 |
| Vector distances | **0.80** | 0.94 | **0.69** |
| R-index | 0.59 | **0.96** | 0.42 |
| DTW | 0.75 | 0.94 | 0.62 |
| *H. sapiens* |  |  |  |
| Hash-based | 0.66 | 0.87 | 0.53 |
| FM-index | 0.01 | 0.05 | 0.01 |
| Vector distances | 0.26 | 0.57 | 0.16 |
| R-index | 0.66 | 0.85 | 0.54 |
| DTW | **0.75** | **0.94** | **0.62** |

**Table 5:** Read classification quality (F1, Precision, Recall) using different matching techniques.

| *Zymo* |  |  |  |
| --- | --- | --- | --- |
| Matching Method | F1 | Precision | Recall |
| Hash-based | 0.95 | 0.92 | 0.97 |
| FM-index | 0.62 | 0.45 | **0.99** |
| Vector distances | 0.96 | 0.97 | 0.95 |
| R-index | 0.96 | **1.0** | 0.93 |
| DTW | **0.98** | 0.99 | 0.97 |

Second, vector distance-based matching shows exceptional quality on simpler genomes but suffers from dramatic quality degradation with increasing genomic complexity. It achieves a high F1 score (0.83) for *E. coli* mapping, comparable to hash-based approach, but quality drops to 0.67 for *D. melanogaster* and 0.26 for *H. sapiens*. This decline suggests poor scaling with increased noise and repetitive content characteristics of more complex genomes.

Third, FM-index demonstrates poor quality in read mapping with low F1 scores across all organisms. Consistently high precision but extremely low recall likely reflects the mismatch between exact string matching logic and the continuous nature of raw signals.

Fourth, mapping quality consistently reduces from *E. coli* to *H. sapiens* across most methods, with the exception of FM-index which maintains uniformly poor quality. Hash-based methods show the most graceful degradation, while vector distances exhibit the steepest decline. This trend underscores the scalability challenges in RSA, where increased genome complexity, repetitive content, and heterozygosity create increasingly difficult matching problems that current methods struggle to address effectively.

Fifth, the classification results (see Table 5) reveal a different quality landscape in terms of matching method effectiveness. R-index achieves perfect precision and the highest F1 score for classification, while vector distances also excel with an F1 score of 0.95. This trend indicates that while these methods struggle with fine-grained mapping, they are highly effective at organism-level discrimination tasks where approximate matches may be sufficient.

Together, these results underscore that mapping and classification favor different approaches: the former demands discriminative, fine-grained alignment, while the latter benefits from relaxed approximate matching. This suggests that future developments in matching algorithms should consider task-specific optimizations.

#### Assisting Basecalled Analysis

Table 6 shows coverage results comparing RSA-assisted versus RSA-unassisted basecalling across two organisms. We make two key observations.

**Table 6:** Basecalled read mapping quality analysis.

| Dataset | RSA Pre-filter | Average Depth of Cov. (×) | Breadth of Coverage (%) | Aligned Reads (#) |
| --- | --- | --- | --- | --- |
| *E. coli* | ✓<br>✗ | 164.39<br>**184.04** | 82.42<br>**82.49** | 182,871<br>**221,651** |
| *D. melanogaster* | ✓<br>✗ | 11.61<br>**15.24** | 99.82<br>**99.91** | 152,601<br>**251,868** |

First, RSA assistance lowers average depth of coverage while keeping breadth nearly unchanged. In E. coli, depth drops from 184.04× to 164.39×, yet breadth stays at 82.4%. In D. melanogaster, depth decreases from 15.24× to 11.61× with breadth remaining 99.9%. This trend shows that RSA filtering reduces redundant coverage and basecalling while maintaining completeness.

Second, genome complexity determines the effectiveness of RSA assistance. *D. melanogaster* maintains nearly identical breadth coverage despite 39% fewer reads (99.82% vs 99.91%) with RSA assistance which suggests that lower-quality reads are filtered out, yielding sensitive but fewer mappings. *E. coli* shows more modest improvements with 17% fewer reads and minimal breadth change (82.42% vs 82.49%), suggesting that RSA assistance is more beneficial with complex genomes.

These results demonstrate that RSA assistance in basecalling provides substantial benefits for complex genomes by substantially reducing the basecalling load. Although we exclude *H. sapiens* due to performance overheads incurred by the RSA pre-filtering, results suggest that tailored RSA tools could be developed to enable RSA assistance benefits for organisms with larger genomes, potentially across the full spectrum of genomic complexity.

Next, we examine the effect of limited context length on basecallers to evaluate their benefits in real-time analysis where a portion of raw signals is basecalled rather than the entire raw signal. Table 7 shows the quality of Dorado SUP model with varying context lengths in terms of number of chunks of raw signal points across three organisms. We make two key observations.

**Table 7:** Effect of limited number of chunks on Dorado (SUP) basecaller.

| Dataset | Chunks (#) | TP | FP | NA |
| --- | --- | --- | --- | --- |
| *E. coli* | 1 | 84.50 | 2.90 | 12.60 |
|  | 2 | 84.99 | 2.54 | 12.46 |
|  | 5 | 85.77 | 2.15 | 12.07 |
|  | 10 | **88.47** | **1.24** | **10.29** |
| *D. melanogaster* | 1 | 78.30 | 20.40 | 1.30 |
|  | 2 | 83.46 | 15.40 | 1.14 |
|  | 5 | 89.85 | 9.23 | 0.92 |
|  | 10 | **94.10** | **5.31** | **0.59** |
| *H. sapiens* | 1 | 80.67 | 10.80 | 8.52 |
|  | 2 | 80.56 | 11.12 | **8.32** |
|  | 5 | 81.89 | 9.50 | 8.61 |
|  | 10 | **84.07** | **7.32** | 8.61 |

First, genome complexity determines sensitivity to context length limitations. *D. melanogaster* demonstrates the strongest response to increased context length, with TP rates improving from 78.30% to 94.10% and FP rates decreasing from 20.40% to 5.31% as context length increases. In contrast, *E. coli* shows modest improvements while *H. sapiens* exhibits the least amount of gains. These results indicate that additional context enables better basecalling decisions, but its benefits diminish for *H. sapiens* where extra context may not resolve ambiguities in homopolymeric regions.

Second, Read Until [62] applications can achieve substantial quality gains with relatively modest increases in context length, particularly from 1 to 5 chunks. Largest benefits occur in the first few context length increases, with diminishing returns beyond the first 5 chunks. This finding indicates that Read Until strategies could strike a balance between performance and quality by using adaptive context lengths, especially given that *D. melanogaster* achieves 89.85% TP with 5 chunks compared to 94.10% with 10 chunks.

The findings suggest that adaptive context length strategies, tailored to the expected genomic complexity of the target organism, could optimize Read Until sequencing quality while managing computational overhead. The substantial improvements observed with even modest increases in context length (from 1 to 2-5 chunks) indicate that small increases in sequencing time can yield significant gains in basecalling quality, for some organisms.

### 5.3 Performance Evaluation

#### Pore Models

Table 8 summarizes the read mapping performance of different pore models. We make two key observations.

**Table 8:** Read mapping performance using different pore models.

| Pore Model | Elapsed time<br>(hh:mm:ss) | CPU time<br>(sec) | Peak<br>Mem. (GB) |
| --- | --- | --- | --- |
| *E. coli* |  |  |  |
| ONT | 0:08:22 | 6,481 | 4.46 |
| Uncalled4 | **0:05:51** | **5,730** | **4.36** |
| *D. melanogaster* |  |  |  |
| ONT | 2:41:07 | 596,614 | 10.25 |
| Uncalled4 | **2:07:02** | **462,608** | **9.6** |
| *H. sapiens* |  |  |  |
| ONT | 1:46:47 | 189,846 | **80.02** |
| Uncalled4 | **0:53:03** | **186,301** | 91.96 |

First, Uncalled4 provides substantial runtime benefits for smaller and intermediate genomes. For *E. coli* and *D. melanogaster*, Uncalled4 reduces elapsed time by 20–50% relative to ONT, while also reducing CPU time and memory footprint. These improvements suggest that the more compact Uncalled4 pore representation accelerates signal processing without sacrificing quality, consistent with the quality results in Table 2.

Second, performance benefits of Uncalled4 diminish for larger genomes. For *H. sapiens*, Uncalled4 enables faster execution with a higher peak memory demand. This indicates a shift in bottlenecks, where reduced computational overhead is offset by increased memory usage, again likely due to the denser signal lookup tables created using Uncalled4 when applied to complex genomes [84].

These results highlight the importance of aligning pore model choice with both dataset scale and available system resources.

#### Segmentation Methods

Tables S4 and S5 show the performance of different segmentation methods. We note one key pattern. Move tables, precomputed by the Dorado (SUP) basecaller, are supplied externally to the RSA pipeline. To ensure fairness, we report results including the computation time for move tables. In this setting, move tables come with additional GPU runtime, while offering no improvement over the simpler t-test segmentation. This suggests that move tables are not yet optimized for RSA. However, targeted refinement of move tables for raw signal segmentation, similar to their utility in other contexts [85, 88, 108], could enable them to outperform current methods in future iterations.

#### Matching Methods

Table 9 summarizes the performance of different matching methods. We make three key observations.

**Table 9:** Read mapping performance using different matching methods.

| Matching Method | Elapsed time<br>(hh:mm:ss) | CPU time<br>(sec) | Peak<br>Mem. (GB) |
| --- | --- | --- | --- |
| *E. coli* |  |  |  |
| Hash-based | 0:05:51 | 5,730 | 4.36 |
| FM-index | 6:57:45 | 1,603,653 | **1.09** |
| Vector distances | 0:20:10 | 54,310 | 54.32 |
| R-index | **0:04:25** | **4,224** | 1.4 |
| DTW | 0:09:23 | 6,128 | 4.43 |
| *D. melanogaster* |  |  |  |
| Hash-based | 2:07:02 | 462,608 | 9.6 |
| FM-index | 3:56:13 | 892,824 | **1.49** |
| Vector distances | 3:22:15 | 823,117 | 255.97 |
| R-index | 1:22:30 | 310,695 | 3.1 |
| DTW | **0:24:02** | **88,044** | 10.46 |
| *H. sapiens* |  |  |  |
| Hash-based | 0:53:03 | 186,301 | 91.96 |
| FM-index | **0:08:44** | **32,808** | **7.52** |
| Vector distances | 5:59:29 | 1,238,190 | 265.16 |
| R-index | 0:35:02 | 131,095 | 29.43 |
| DTW | 0:46:35 | 158,289 | 116.2 |

First, R-index offers a balance between speed and memory footprint. Across all organisms, R-index delivers the fastest execution while keeping memory demands low. This trend favors the use of R-index for real-time and large-scale analyses.

Second, vector distance methods are computationally demanding. While achieving strong quality (Table 4), they require orders of magnitude more memory and runtime—up to six hours and 265 GB for the *H. sapiens* dataset. This resource intensity makes FM-index a practical alternative for complex genomes despite the headroom for quality.

Third, hash-based and DTW matching occupy intermediate positions with distinct trade-offs. Hash-based matching is more memory-efficient, while DTW, by contrast, requires more memory but substantially less than vector distances, with runtimes that scale gracefully with genome size.

The results highlight that no single method dominates: R-index is fastest on *E. coli*, DTW on *D. melanogaster*, and FM-index leads on *H. sapiens* and in peak memory demand across organisms. Vector distances methods are consistently the most memory-intensive. The changing trends indicate that method choice should be made based on genome complexity and resource limitations.

## 6 Resources

The RawBench framework, including datasets, evaluation scripts, and documentation, is available on https://github.com/CMU-SAFARI/RawBench. The framework is publicly available to facilitate its adoption and extension by the research community.

## 7 Conclusion

RawBench provides a comprehensive benchmarking framework for raw signal analysis (RSA) that addresses critical gaps in current frameworks. By decomposing analysis pipelines into three modular components (i.e., reference genome encoding, signal encoding, and representation matching) and evaluating them across organisms of varying genomic complexity, we demonstrate that statistical segmentation methods outperform the intermediate outputs of basecallers, though carefully designed context-aware ML models can surpass these statistical approaches. We further show that advanced pore models like Uncalled4 offer consistent quality improvements and hash-based matching provides the most robust quality across genome complexities. The framework’s modular design enables systematic evaluation of emerging methods while maintaining biological relevance via up-to-date datasets. RawBench establishes a foundation for systematic progress in RSA, enabling researchers to design more effective pipelines and unlock the full potential of raw signal data for diverse genomics applications.

## Acknowledgments

We thank the anonymous reviewers of ACM BCB 2025 for their valuable feedback. We thank the SAFARI Research Group members for thoughtful feedback and the stimulating intellectual & scientific environment they cultivate. We acknowledge the generous support of our industrial partners, including Google, Huawei, Intel, and Microsoft. This work is partially supported by the ETH Future Computing Laboratory (EFCL) and the European Union’s Horizon research and innovation programme [101047160 BioPIM].

## Supplementary Material

## A Extended Quality Benchmarks

**Table S1:** Read classification quality (F1, Precision, Recall) using different pore models.

| *Zymo* |  |  |  |
| --- | --- | --- | --- |
| Pore Model | F1 | Precision | Recall |
| ONT | 0.96 | 1.0 | 0.93 |
| Uncalled4 | 0.96 | 1.0 | 0.93 |

**Table S2:**
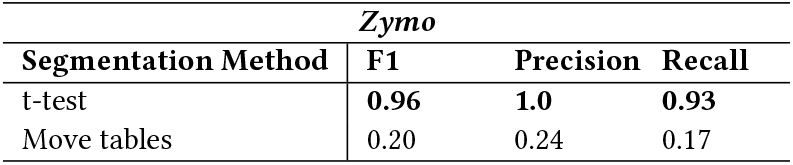
Read classification quality (F1, Precision, Recall) for different segmentation methods.

## B Extended Performance Benchmarks

**Note**. *NA* in Tables S4 and S5 indicates the metric was not applicable for that run.

**Table S3:** Read classification performance using different pore models.

| Pore Model | Elapsed time<br>(hh:mm:ss) | CPU time<br>(sec) | Peak<br>Mem. (GB) |
| --- | --- | --- | --- |
| *Zymo* |  |  |  |
| ONT | 0:03:42 | 792 | 2.53 |
| Uncalled4 | 0:04:31 | 1,085 | 2.53 |

**Table S4:** Read mapping performance using different segmentation methods.

| Segmentation Method | Elapsed time<br>(hh:mm:ss) | CPU time<br>(sec) | Peak<br>Mem. (GB) |
| --- | --- | --- | --- |
| *E. coli* |  |  |  |
| t-test | 0:05:51 | 5,730 | 4.36 |
| Move tables | 0:38:52 | NA | 14.64 |
| *D. melanogaster* |  |  |  |
| t-test | 2:07:02 | 462,608 | 9.6 |
| Move tables | 1:39:47 | NA | 18.25 |
| *H. sapiens* |  |  |  |
| t-test | 0:53:03 | 186,301 | 91.96 |
| Move tables | 0:53:26 | NA | 93.18 |

**Table S5:** Read classification performance using different segmentation methods.

| Segmentation Method | Elapsed time<br>(hh:mm:ss) | CPU time<br>(sec) | Peak<br>Mem. (GB) |
| --- | --- | --- | --- |
| *Zymo* |  |  |  |
| t-test | 0:04:31 | 1,085 | 2.53 |
| Move tables | 0:12:57 | NA | 3.95 |

**Table S6:**
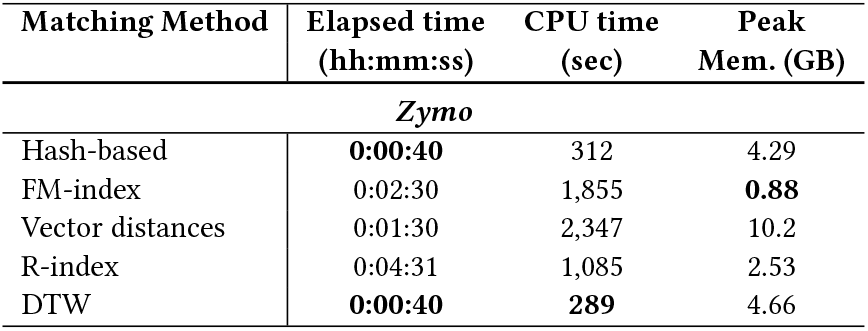
Read classification performance using different matching methods.

| Matching Method | Elapsed time<br>(hh:mm:ss) | CPU time<br>(sec) | Peak<br>Mem. (GB) |
| --- | --- | --- | --- |
| *Zymo* |  |  |  |
| Hash-based | 0:00:40 | 312 | 4.29 |
| FM-index | 0:02:30 | 1,855 | 0.88 |
| Vector distances | 0:01:30 | 2,347 | 10.2 |
| R-index | 0:04:31 | 1,085 | 2.53 |
| DTW | 0:00:40 | 289 | 4.66 |

## C Configuration

### C.1 Parameters

In Supplementary Table S7, we show the details of the parameters used for each tool and dataset including preset values, when required. For minimap2 [103], we use the same parameter setting for all datasets. For the Dorado super-accurate (SUP) basecaller, we use the model trained for the corresponding data sampling frequency (i.e. 4 kHz or 5 kHz). Thread count was specified as 64 for all CPU workloads, i.e., all processes other than basecalling which used a GPU.

**Table S7:**
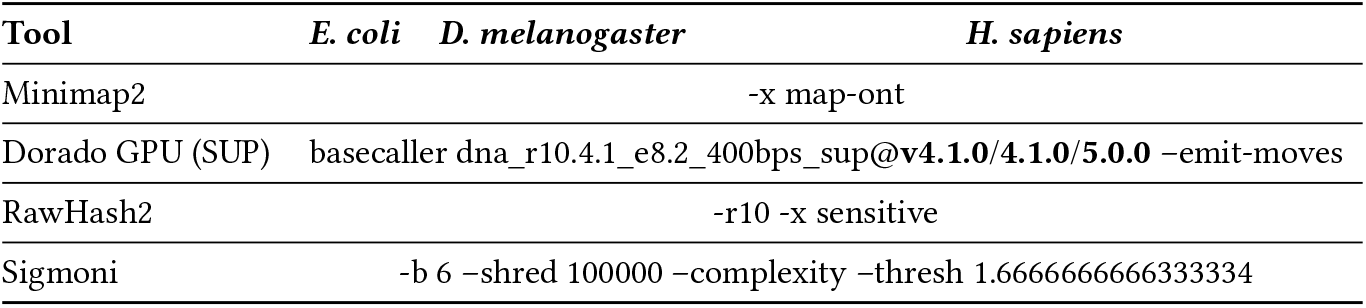
Parameters we use in our evaluation for each tool and dataset in mapping. Tool *E. coli D. melanogaster*.

### C.2 Versions

Supplementary Table S8 shows the version and the link to these corresponding versions of each tool we use in our experiments. Scripts to reproduce each of the experiments can be found on https://github.com/CMU-SAFARI/RawBench.

**Table S8:** Versions of each tool and library.

| Tool | Version | Link to the Source Code |
| --- | --- | --- |
| RawHash2 | 2.1 | https://github.com/CMU-SAFARI/RawHash/releases/tag/v2.1 |
| Minimap2 | 2.28-r1209 | https://github.com/lh3/minimap2/releases/tag/v2.28 |
| Dorado | 0.9.6 | https://github.com/nanoporetech/dorado/releases/tag/v0.9.6 |
| Sigmoni |  | https://github.com/vikshiv/sigmoni |
| UNCALLED | 2.3 | https://github.com/skovaka/UNCALLED/releases/tag/v2.3 |
| Uncalled4 | 4.1.0 | https://github.com/skovaka/uncalled4/releases/tag/4.1.0 |
| Sigmap | 0.1 | https://github.com/haowenz/sigmap/releases/tag/v0.1 |
| Mosdepth | 0.3.10 | https://github.com/brentp/mosdepth/releases/tag/v0.3.10 |
| samtools | 1.22 | https://github.com/samtools/samtools/releases/tag/1.22 |
| bedtools | 2.31.1 | https://github.com/arq5x/bedtools2/releases/tag/v2.31.1 |

## Notes

### Competing Interest Statement

The authors have declared no competing interest.

